# *foxc1a* genetically interacts with *ripply1* to regulate *mesp-ba* expression and somitogenesis in the zebrafish embryo

**DOI:** 10.1101/041012

**Authors:** Rotem Lavy, W. Ted Allison, Fred B. Berry

## Abstract

Somitogenesis is a fundamental segmentation process that forms the vertebrate body plan. A network of transcription factors is essential in establishing the spatial temporal order of this process. One such transcription factor is mesp*-ba* which has an important role in determining somite boundary formation. Its expression in somitogenesis is tightly regulated by the transcriptional activator Tbx6 and the repressor Ripply1 via a feedback regulatory network. Loss of *foxc1a* function in zebrafish leads to lack of anterior somite formation and reduced *mesp-ba* expression. Here we examine how *foxc1a* interacts with the *tbx6-ripply1* network to regulate *mesp-ba* expression. In *foxc1a* morphants, anterior somites did not form at 12.5 hours post fertilization (hpf). At 22 hpf posterior somites formed, whereas anterior somites remained absent. In *ripply1* morphants, no somites were observed at any time point. The expression of *mesp-ba* was reduced in the *foxc1a* morphants and expanded anteriorly in *ripply1* morphants. The *tbx6* expression domain was smaller and shifted anteriorly in the *foxc1a* morphants. Double knockdown of *foxc1a* and *ripply1* resulted in absence of anterior somite formation while posterior somites did form, suggesting a partial rescue of the *ripply1* phenotype. However, unlike the single *foxc1a* morphants, expression of *mesp-ba* was restored in the anterior PSM. Expression of *tbx6* was expanded anteriorly in the double morphants. In conclusion, both *foxc1a* and *ripply1* morphants displayed defects in somitogenesis, but their individual loss of function had opposing effects on *mesp-ba* expression. Loss of *ripply1* appears to have rescued the *mesp-*ba expression in the *foxc1a* morphant, suggesting that intersection of these parallel regulatory mechanisms is required for normal *mesp-ba* expression and somite formation.

## 1. Introduction

Somitogenesis is a complex segmentation process common to all vertebrate species. Multiple regulatory pathways and networks drive the formation of somites early in embryonic development. In mammals, the somites contain precursors for the bony components of the axial skeleton, as well as the muscle and skin of the trunk [1–4]. Defects in somite formation lead to abnormal development of the skeleton and can manifest in fused ribs and vertebrae, pebble-like vertebral formation and scoliosis [5]. In zebrafish, the somites contain predominantly myotome precursors required for the development of strong trunk muscle needed for locomotion in an aqueous environment, with sclerotomal precursors a minor component of the somites [6]. Nonetheless, defects in somite segmentation in zebrafish have been shown to result in vertebral patterning malformations leading to scoliotic phenotypes [7].

While the mechanisms driving somitogenesis are still not completely understood, one of the main models describing it is the clock and wavefront segmentation model, which postulates that cyclic waves of gene expression sweep through the presomitic mesoderm (PSM) in a posterior to anterior manner until they reach a boundary determination front in the anterior PSM [1, 8–12]. The cells in the anterior PSM, where the wavefront meets the boundary determination front, must be in a permissive state for subsequent differentiation and mesenchymal to epithelial transformation, allowing the formation of intersomitic boundaries. The spatial and temporal expression of the regulatory components involved in somitogenesis is thus thought to be crucial for proper somite formation and subsequent skeletal development.

The transcription factor mesoderm posterior protein 2 (MESP2) is indispensable for somitogenesis. *Mesp2* regulates boundary formation and rostro-caudal patterning of the somite [13]. Mutations in *Mesp2* in mice leads to loss of intersomitic boundary formation, fused somites, and subsequently fused ribs and vertebrae [13, 14]. In humans, *MESP2* mutations are the underlying cause of spondylocostal dysostosis, a condition characterized by misshapen and/or fused ribs and vertebrae and scoliosis [15]. The spatial expression of *Mesp2* is regulated by the Notch pathway, whereas its temporal expression is regulated via the Tbx6-Ripply regulatory network in a negative feedback loop in which Tbx6 induces *Mesp2* expression. As levels of Mesp2 increase expression of *Ripply* is induced. Ripply then binds to TBX6 converting this complex to a transcriptional repressor as well as promoting the degradation of TBX6 and thus preventing further *Mesp2* induction [13, 16]. *Mesp2* expression is not detectable in mouse and zebrafish embryos lacking Tbx6, and somite segmentation was severely defective [17, 18]. Ripply1 and Ripply2 have been shown to interact with Tbx6 to regulate the temporal expression of *Mesp2* [3, 19]. In the absence of Ripply2 in mice, or its homologue ripply1 in zebrafish, no segmental borders are detected throughout somitogenesis, and the expression of *Mesp2* and *Tbx6* are upregulated and extend anteriorly in PSM [3, 19].

FOXC1, a member of the <u>Fo</u>rkhead Bo<u>x</u> transcription factor family, also has a crucial, dose dependent role in somitogenesis. Mice with compound homozygous mutations in *Foxc1* and its paralogue *Foxc2* have a complete lack of somite formation [20]. The expression of *Mesp2* is strongly downregulated in the PSM in these embryos. Mice with homozygous null-Foxc1 mutations have severe axial skeletal defects such as incomplete formation of the neural arches, reduced centra of the vertebrae, and fused ribs as well as anomalies in other developmental processes, such as ocular, cardiovascular and genitourinary abnormalities [21].

In zebrafish there are two copies of *foxc1*, *foxc1a* and *foxc1b*, whereas *Foxc2* ortholog have not been found. Foxc1a seems to play a role in somitogenesis, as double knockdowns of both genes produced similar results to those of Foxc1a single knockdowns [22]. When *foxc1a* was knocked down in zebrafish, formation of the anterior somites was defective, and the spatial polarity of the PSM was lost. Additionally, the expression of *mesp-ba*, a zebrafish *Mesp2* homologue, was absent in *foxc1a* morphants, mimicking the mouse phenotype, and suggesting that Foxc1 has a conserved role in the regulation of *Mesp2* expression [22].

Here we sought to determine the mechanism through which Foxc1a regulates the expression of *mesp-ba* and the segmentation of the somite. We show that *foxc1a* genetically interacts with *ripply1* to regulate *mesp-ba* and *tbx6* expression, possibly through an interaction with Fgf8.

## 2. Results

To analyze the role of Foxc1a in somite formation, we have used a well-characterized antisense morpholino-modified oligonucleotide (MOs) targeting the mRNA translation start site [22]. As previously shown, injections with Foxc1a MO resulted in loss of somite formation at 12.5 hours post fertilization (hpf), compared with the control-injected embryos which were at the 3-somite stage at this time point (Fig 1A, B) as well as at 14 hpf, when control embryos were at the 9-somite stage (Fig 1A’, B’). At 22 hpf posterior somites did form in both the *foxc1a* morphants and the control injected embryos (Fig 1A”, B”). To validate the specificity of the knockdown effect seen, zebrafish embryos were co-injected with Foxc1a MO and 20pg *foxc1a* mRNA resistant to the morpholino. Somite formation in these embryos proceeded normally and at a rate comparable to that of the control-injected embryos (Fig1 E-G).

**Fig 1.**
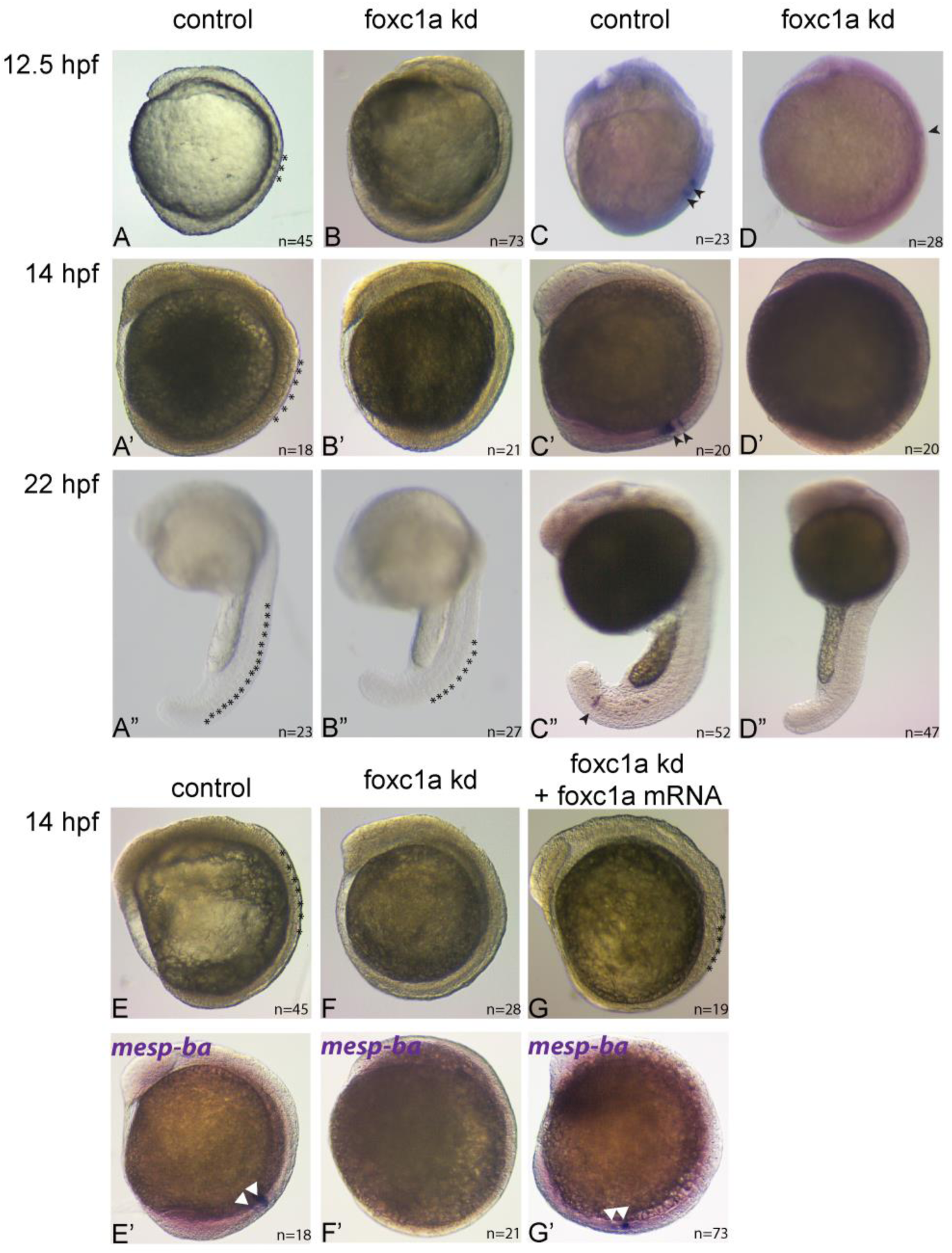
Foxc1a knockdown prevents anterior somite formation and reduces *mesp-ba* expression throughout somitogenesis. Phase contrast micrographs of embryos injected with control morpholinos (A, B, C) and *foxc1a* morpholinos (A’,B’,C’) at 12.5, 14 and 22 hpf, respectively. Somites are indicated with asterisks. Whole mount in situ hybridization of *mesp-ba* mRNA localization on embryos at 12.5 hpf (D, D’), 14 hpf (E, E’) and 22 hpf (F, F’) injected with control morpholinos (D-F) or foxc1a morpholinos (D’-F’). Arrowheads denote the localization of mesp-ba expression domains. Phase contrast micrographs (E-G) and whole mount in-situ hybridization of *mesp-ba* mRNA localization (E’-G’) of embryos injected with control morpholinos (E, E’), *foxc1a* morpholinos (F, F’) or *foxc1a* morpholino and *foxc1a* mRNA (G, G’).

The expression of *mesp-ba* in our control embryos was detected in 1-2 bands in the anterior PSM throughout somitogenesis (12.5, 14, and 22 hpf, Fig 1C-C”). In the *foxc1a-*morphants, *mesp-ba* expression was reduced in the anterior PSM at all time points tested (Fig 1D-D”). When *foxc1a* mRNA was injected in addition to the MO, *mesp-ba* expression was rescued that to the level seen in the control embryos (Fig 1 F’-G’).

Next, we examined whether Foxc1 interacts with the Tbx6-Ripply network to regulate *mesp-ba* expression. First we looked at whether *tbx6* expression was affected in the *foxc1a* morphants. In control embryos, *tbx6* was detected throughout the PSM, but was not detected in the tailbud. In *foxc1a* morphants, while *tbx6* was detected, its expression domain was smaller and more anteriorly shifted compared with control embryos (Fig 2). As both *foxc1a* and *ripply1* morphant fish are characterized by an absence of posterior somites we postulated that these factors may genetically interact to regulate the expression of *tbx6* and its downstream target *mesp-ba*. We knocked down the expression of *ripply1* using a previously characterized translation-blocking morpholino [3], and observed a complete lack of segmentation both at 12.5 hpf and 22 hpf (Fig 3A”, B”). The expression domains of both *tbx6* and *mesp-ba* were extended anteriorly into the somitic mesoderm (Fig 3). Interestingly, when the *ripply1* and *foxc1a* MOs were co-injected, anterior somites failed to form (Fig 3A’”), while posterior somite did form (Fig 3B’”), resulting in a similar morphological phenotype as observed in the Foxc1a single morphants (Fig 3A’, B’, respectively). However, unlike single Foxc1a morphants, *mesp-ba* mRNA expression was detected in 1-2 bands in the anterior PSM (Fig 3C, D, C’”, D’”), and the *tbx6* expression domain was also restored in the PSM (Fig 3E, E’”, F, F’”).

**Fig 2.**
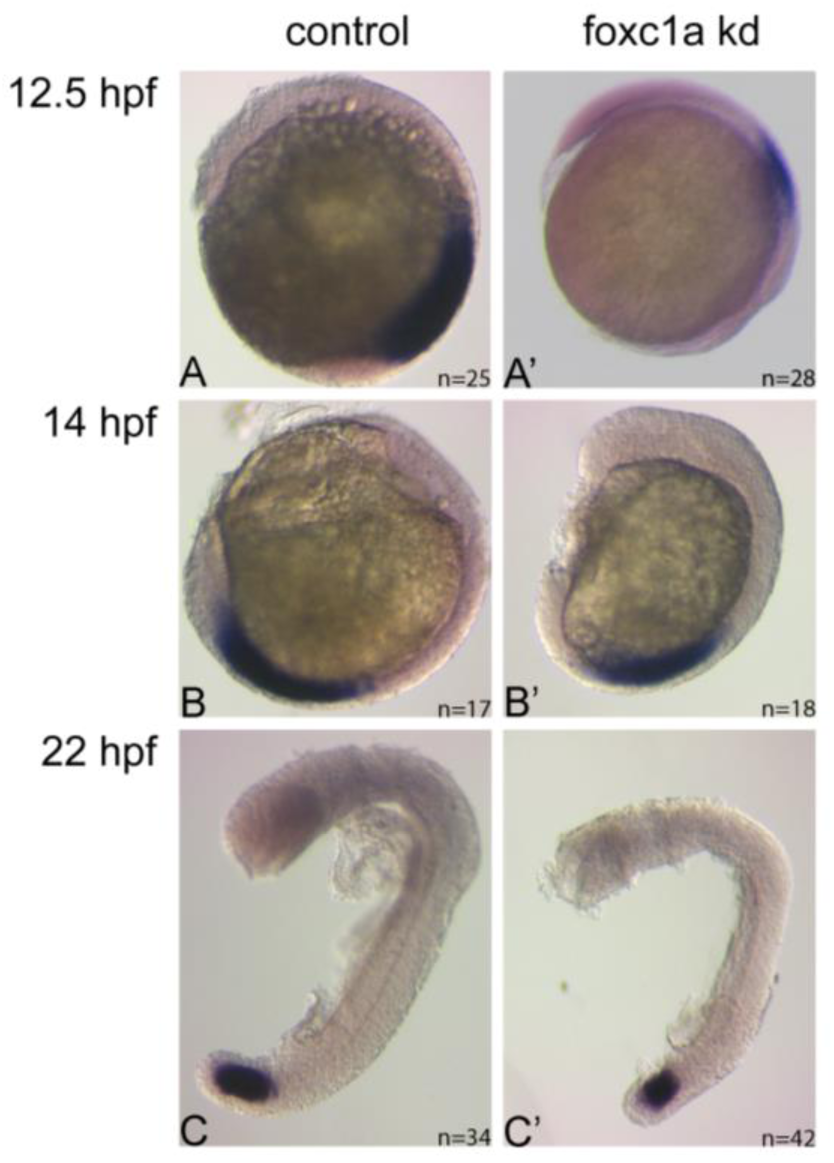
The *tbx6* expression domain is smaller and anteriorly shifted when *foxc1a* is knocked down. Whole mount in-situ hybridization of *tbx6* mRNA DIG-labeled probe of control-(A, B, C) or *foxc1a*-MO (A’, B’, C’) injected embryos at 12.5, 14, and 22 hpf, respectively.

**Fig 3.**
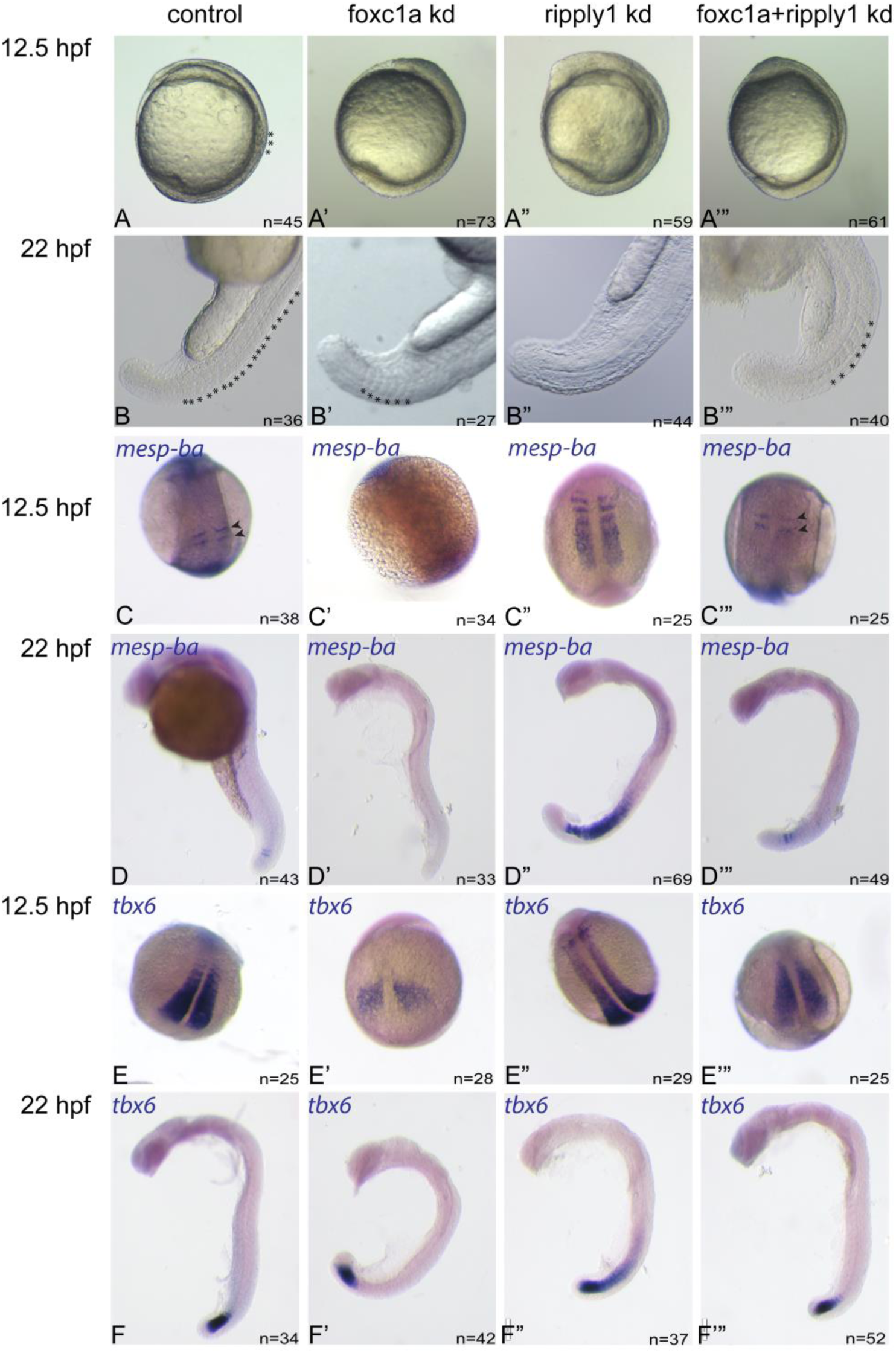
*ripply1* MO rescues altered *mesp-ba* and *tbx6* expression observed in *foxc1a* morphants. Embryos injected with control, *foxc1a*, *ripply1* or *foxc1a+ripply1* morpholino(s) (A-F, A’-F’, A”-F”, and A’”-F’”, respectively). Phase contrast imaging (A-A’”, B-B’”) at 12.5 and 22 hpf. Somites are indicated with asterisks. Whole mount in-situ hybridization of *mesp-ba* mRNA localization (C-C’”, D-D’”) at 12.5 or 22 hpf, respectively. *mesp-ba* expression domain is indicated with arrowheads. Whole mount in-situ hybridization of *tbx6* mRNA localization (E-E’”, F-F’”) at 12.5 or 22 hpf, respectively.

We also wanted to determine whether the Fgf8 gradient is affected in our morphant embryos. The expression of *mesp-ba* has been shown to be affected by the level of Fgf8a in the PSM [26], and we speculated that Foxc1a might exert its effect on *mesp-ba* through Fgf8. We therefore examined the expression of *fgf8a* in our control, *foxc1a*-, *ripply1-*, and *foxc1a-*ripply1-morphants at 14 hpf. In our control-injected embryos *fgf8a* was detected in 2-3 stripes in the midbrain-hindbrain boundary, in the tailbud, in the anterior PSM and in the anterior domain of the newly formed somites (Fig 4A). In the *foxc1a* morphants, *fgf8a* expression was detected in the tailbud and in the midbrain-hindbrain boundary, but was reduced in the PSM and in the area where normally somites would have formed (Fig 4B). In *ripply1* morphants, *fgf8a* expression resembled that of the control-injected embryos and was detected in the tailbud, midbrain-hindbrain boundary, and the PSM. While these morphants don’t have clearly segmented somites, *fgf8a* expression in the PSM seemed to span the same area it did in the control embryos (Fig 4C). The expression of *fgf8a* in the *foxc1a-ripply1* double morphants also resembled that of the control embryos, however its expression in the PSM appeared to be reduced compared to the controls and *ripply1*-morphants (Fig 4D).

**Fig 4.**
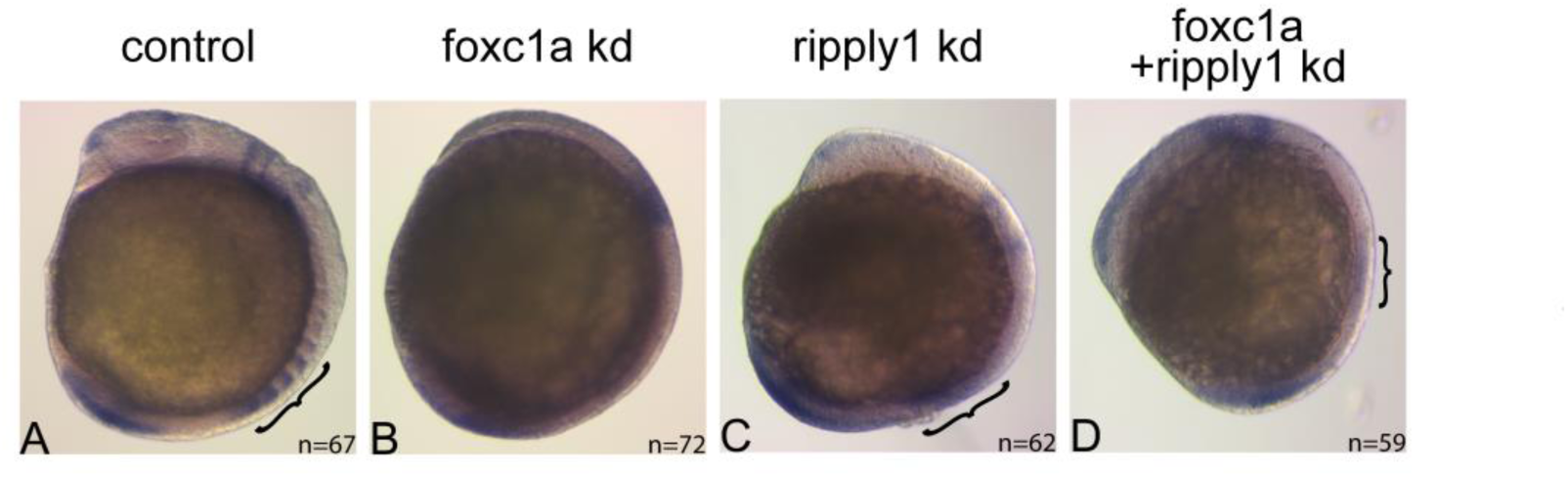
*ripply1* MO rescues the expression of *fgf8a* in the PSM in the absence of *foxc1a*. Embryos injected with control (A), *foxc1a* (B), *ripply1* (C) or *foxc1a+ripply1* morpholino(s) (D). Phase contrast imaging of whole mount in-situ hybridization of *fgf8a* mRNA localization at 14 hpf. Brackets denote expression in the PSM.

## 3. Discussion

Accumulating evidence in recent years suggests that Foxc1 has an important, dose dependent role in somite formation and axial skeletal development [20, 22–24]. Mice with null Foxc1 mutations have defects in axial skeletal development, such as incomplete formation of the neural arches, misshapen ribs, and fused vertebrae, along with cardiovascular, ocular and other anomalies [20]. In zebrafish, it has been demonstrated that disruption of Foxc1a function results in lack of somite formation at the 7-somite stage [22]. More recently, it has been demonstrated that in *foxc1a* mutants anterior somite formation was altered [23]. However, how *foxc1a* exerts its function on somitogenesis is still fully not understood. We show here that *foxc1a* genetically interacts with *ripply1* to regulate *mesp-ba* expression and somite formation. To discern how Foxc1 functions during somitogenesis and to assess the mechanism through which it acts to regulate *mesp-ba* expression, we have utilized Morpholino-mediated gene knockdowns. *Foxc1a*-morpholinos have been used in the past to learn of its effect on several developmental processes, including angiogenesis and cardiac development [25–28]. When we knocked *foxc1a* down in zebrafish embryos, we observed a lack of anterior somite formation, whereas posterior somites continued to be normally formed. This is likely not the result of the morpholino having transient effects, since down regulation of *mesp-ba* expression persisted in these later forming somites. When we co-injected *foxc1a-*MO with *foxc1a* mRNA these phenotypes were rescued, suggesting that the phenotypes observed are specific and result from reduced Foxc1a expression.

Mammals possess two copies of *Foxc* genes, *Foxc1* and *Foxc2*. Zebrafish, on the other hand, do not possess a *Foxc2* orthologue but have 2 homologues of *foxc1*: *foxc1a* and *foxc1b*. While both *Foxc1* and *Foxc2* are expressed in the presomitic tissue in mice, the phenotypes observed in the compound, homozygous *Foxc1^-/-^ Foxc2^-/-^* mice are far more severe than those observed in the single *Foxc1*-null or *Foxc2*-null mice, suggesting that the two gene do share some compensatory roles. In zebrafish, it appears that *foxc1b* does not regulate somitogenesis, and the phenotypes observed when *foxc1a* and *foxc1b* are knocked down together closely resemble those of a single *foxc1a* knockdown [22]. We have thus decided to focus on the role of *foxc1a* in somitogenesis.

In addition to the lack of anterior somite formation when *foxc1a* was knocked down, the expression of *mesp-ba* was reduced throughout somitogenesis, and the expression domain of *tbx6* was shifted anteriorly and appeared smaller. When *ripply1*, a component of the mesp-ba–tbx6 regulatory network was knocked down, a complete lack of somite formation was observed. The expression domains of both *mesp-ba* and *tbx6* were expanded anteriorly, probably due to lack of repression resulting from the absence of Ripply1. Interestingly, when *foxc1a* and *ripply1* were knocked down together, the morphological phenotype resembled that of the *foxc1a* single morphants and restored formation of the posterior somites was observed. The expression of *mesp-ba* and *tbx6* was also rescued, and resembled those observed in the control embryos. Together these observations suggest that *foxc1a* may be required for the *ripply1* morphant phenotype.

It has been previously shown that the expression of *Mesp2* is dependent on the positioning of the determination front, as defined by the expression of Fgf8 [26]. Additionally, *Foxc1* and *Fgf8* genetically interact during mammalian jaw patterning and *Fgf8* expression is reduced in the absence of *Foxc1* in mice [29]. We therefore decided to examine the expression of *fgf8a* in our morphant embryos. We found that *fgf8a* expression was indeed reduced in the PSM in our *foxc1a* morphants compared to controls. In *ripply1* morphants *fgf8a* expression was still expressed in the anterior PSM even though somites did not form. These data suggest that the lack of *fgf8a* expression in the anterior PSM of *foxc1a* morphants is not due to absence of somite formation and may indicate regulation of *fgf8a* by foxc1a. It is thus possible that the restored expression of *mesp-ba* in the double morphants compared to the *foxc1a* morphants is a result of the rescue of *fgf8a* expression when *ripply1* is absent in addition to *foxc1a*.

These observations suggest that *foxc1a* and *ripply1* genetically interact to regulate the expression of *mesp-ba* in the PSM and somite formation. Our results demonstrate that *foxc1a* has a differential role in somite formation in that anterior somites fail to form in the absence of *foxc1a* but posterior somite segmentation can still occur. This observation is consistent with several recent reports of underlying differences in the formation of the anterior and posterior somites. For example, there is a differential requirement for *Lunatic fringe (Lfng)* expression and Notch oscillations in anterior and posterior body formation [30, 31]. Similarly, mutations in *intergrinα5* disrupt anterior somite formation, while posterior somites are unaffected [32]. Taken together, these observations suggest that the formation of the anterior somites is regulated via different molecular mechanisms than those regulating the formation of the posterior somites, however the mechanistic underpinnings of these surprising differences remain to be understood.

We also demonstrate here that *mesp-ba* can be detected when *foxc1a* is knocked down in combination with and *ripply1*, suggesting that Foxc1a does not directly regulate expression of mesp-ba. Instead, we suggest that Foxc1a regulates the expression of *mesp-ba* through its interaction with *fgf8a*. The absence of *ripply1* in the double morphants rescues the expression of *fgf8a* in the PSM, thus allowing for the specification of the determination front and the expression of *mesp-ba*. Importantly, the detection of *mesp-ba* and *tbx6* in the double morphants when Ripply1 is absent suggests that these genes could be regulated by a mechanism independent of Ripply1. Finally, the formation of the posterior somites in the *foxc1a* single morphants despite the reduced expression of *mesp-ba* demonstrates that segmentation of the somites can occur in the absence of *mesp-ba* expression, and that the expression of *mesp-ba* can be regulated through a mechanism independent of the Tbx6-Ripply1 network, as demonstrated by the restored expression of *mesp-ba* and *tbx6* when both Foxc1a and Ripply1 are lost, compared with the *ripply1* single morphants. Our findings provide additional insight into the progression of somitogenesis and the

## 4. Experimental Procedures

### Animal Ethics Statement

Zebrafish husbandry and experimental procedures complied with the University of Alberta Animal Care and Use Committee guidelines under directives of the Canadian Council on Animal Care.

### Zebrafish and morpholino knockdown

*Danio rerio* were maintained as previously described [33]. Wild-type zebrafish crosses (AB x Wik) were used for this study. Previously characterized translation blocking antisense morpholino oligonucleotides (MOs) against *foxc1a, ripply1* and a standard control were obtained from Gene Tools, LLC (Supplementary Table 1) and injected into embryos at the one to two-cell stage. The *foxc1a* MO was injected at a concentration of 6.5ng/embryo and the *ripply1* MO was injected at a concentration of 3.25ng/embryo. When a single gene was targeted for knockdown the injection solution also contained the standard control MO, such that all embryos were injected with a total MO concentration of 9.75ng/embryo. Embryos were then allowed to grow at 28.5°C. All embryos were euthanized using MS222 and fixed in 4% paraformaldehyde overnight at 4°C.

### mRNA Production and Rescue Injections

Zebrafish *foxc1a* was inserted into a pCS2+ vector and grown in DH5α cells. Plasmid was purified using Qiagen Plasmid *Plus* Midi kit (Qiagen 12943) and linearized using NEB NotI restriction enzyme (R0189S). Linearized DNA was then purified using 0.5M EDTA and 3M sodium acetate pH 5.2 in 100% EtOH and chilled in −80°C overnight. DNA was then pelleted and RNA was transcribed and purified using Ambion mMessage mMachine T7 transcription kit (AM1344). mRNA was aliquoted and stored at −80°C. For rescue experiments, 20 pg mRNA were co-injected with 6.25 ng *foxc1a* MO2 into embryos at the 1-2 cell stage. This injected *foxc1a* mRNA was inert to its cognate MO because the injected mRNA lacked the 5’UTR of the natively expressed *foxc1a* mRNA, and this is principally where this MO binds to affect gene knockdown. Following injections, embryos were grown and euthanized as described above.

### Whole mount in-situ hybridization

Single or double whole mount in-situ hybridization was performed as described in Gongal and Waskiewicz [34]. cDNAs for *foxc1a, ripply1, mesp-ba, tbx6*, and *fgf8a* were used as PCR-based templates for antisense probes synthesis whereby a T7 or T3 binding site was juxtapositioned on the 3’ end of the reverse primer sequence [35] (Supplementary Table 2).

## Acknowledgements

Zebrafish *fgf8a* plasmids were kindly provided by Dr. Andrew Waskiewicz, University of Alberta. F.B.B. is the holder of the Shriners Hospital for Children Chair in Pediatric Scoliosis Research. This research was supported by grants from the Canadian Institutes of Health Research (F.B.B.), the Women and Children’s Health Research Institute (F.B.B.) and from the Natural Sciences and Engineering Research Council of Canada (W.T.A).

